# GenePioneer: A Comprehensive Python Package for Identification of Essential Genes and Modules in Cancer

**DOI:** 10.1101/2024.12.16.628633

**Authors:** Golnaz Taheri, Amirhossein Haerianardakani

## Abstract

**Summary:** We propose a network-based unsupervised learning model to identify essential cancer genes and modules for 12 different cancer types, supported by a Python package for practical application. The model constructs a gene network from frequently mutated genes and biological processes, ranks genes using topological features, and detects critical modules. Evaluation across cancer types confirms its effectiveness in prioritizing cancer-related genes and uncovering relevant modules. The Python package allows users to input gene lists, retrieve rankings, and identify associated modules. This work providing a robust method for gene prioritization and module detection, along with a user-friendly package to support research and clinical decision-making in cancer genomics.

**Availability:** `GenePioneer` is released as an open-source software under the MIT license. The source code is available on GitHub at https://github.com/Golnazthr/ModuleDetection

## Introduction

Cancer arises from alterations in genes, nucleotides, and cellular structures. Somatic cells, mutating an order of magnitude faster than germline cells, are more susceptible to cancer (1). Mutations can modify protein function and alter diverse cellular processes, leading to significant intra- and inter-tumor heterogeneity in biochemistry and histology (2). This heterogeneity complicates cancer treatment and the identification of causative events. The recent focus on mutations underscores the importance of gene-specific analysis to identify cancer driver genes (3). Identifying recurrently mutated genes can aid in predicting cancer progression. However, many cancer-driver genes remain elusive, with many mutations undetected by current methods. Recent studies have revealed novel genes and cancer gene classes (4). A comprehensive catalog of mutations in different frequency range is crucial for identifying dysregulated pathways and potential therapeutic targets (4). However, cancer genes often disrupt a limited number of pathways, particularly those involved in survival, cell division, differentiation, and genome maintenance. Therefore, assessing the pathway-level importance of genes, even those with intermediate or low mutation frequencies, is essential. Despite extensive research into critical genes and cancer-related modules for specific cancer types, a significant knowledge gap persists: most studies focus on isolated cancer types. This specialization limits the application of essential gene identifier models across different cancers (5). Identifying non-specialized genes and modules can provide insights into common patterns, behaviors, similarities, and differences among various cancer types, offering a holistic view of cancer biology.

Despite the growing availability of advanced scientific tools, a comprehensive toolkit for identifying essential genes and cancer-related modules remains unavailable. Such a tool should be rooted in scientific research while remaining accessible to researchers with varying levels of expertise. This underscores the need for an integrated solution that connects advanced bioinformatics techniques with practical applications in cancer genetics. To fill this gap, we introduce `GenePioneer`, a Python package designed to address these challenges.

## Methods

This section proposes a two-step technique for identifying driver genes and modules across 12 different cancer types. In the first step, a network is constructed using biological process terms and a set of mutated genes for each cancer. The Laplacian Score algorithm (LS) is then employed to score genes within this network, and a subset of informative genes with high scores is selected for each cancer (6). In the second step, the MG is used as a heuristic algorithm to identify clusters with the highest scores as candidate modules (8). Significant clusters with high p-values, based on cancer-related pathways, are then identified as a set of cancer-related modules.

### Datasets

For the first step, we collected a list of the most frequent mutations for 12 cancer types from the TCGA dataset (3). We further enriched the TCGA data with additional biological information from the GO dataset (10) from UniProt, a public dataset containing biological processes and their associated genes (9). By incorporating this enriched dataset, we ensure that the proposed Python package is grounded in robust biological data and can uncover meaningful insights into cancer genomics. For evaluating the genes obtained from the first step, we utilized four benchmark datasets CGC (8), Rule (8), HCD (8) and CTAT (8) respectively. These benchmarks serve as a reliable reference set of known cancer-related genes.

### Network Construction

We constructed an undirected weighted graph *G* = (*V, E, W*) as the basis for both gene ranking and module detection methods. In this network, *V* represents the union of two gene sets *V*_1_ and *V*_2_ respectively. The first set, *V*_1_ = {*g*_1_, …, *g*_200_}, comprises the top 200 genes with the highest mutation frequency in each cancer type. To enrich this initial set, we extended it by incorporating additional genes involved in the same biological processes as the original 200 genes, forming *V*_2_ = {*g*_201_, …, *g*_*m*_}. This expansion ensures that the network reflects not only mutational frequency but also the biological context and interactions among genes. Set *E* represents edges in the network and edge *e*_*ij*_, is created between two genes, *g*_*i*_ and *g*_*j*_, if they share at least one common biological process. The way we defined edges in this network ensure us that the network captures both mutation-based relationships and the biological interaction of genes. The weight of the edge *e*_*ij*_, *ω*(*e*_*i*_*e*_*j*_), between two genes *g*_*i*_ and *g*_*j*_ in the network is counted by the number of biological processes in which both genes participate together. We defined five topological features: weight, betweenness, eigenvector, and entropy, to characterize the role and importance of each node within the network (7). Supplementary Table 1 provides detailed information on the network construction for various cancer types. Subsequently, we employed the LS as a feature selection method to rank these features based on their ability to preserve the local structure of the data and rank genes accordingly. Supplementary Table 2 presents the top 5 genes with the highest LS scores for each cancer type.

**Table 1.** The values of our evaluation measure on different benchmarks for ovarian cancer.

| Benchmark | AUC-ROC | Mann-Whitney p-value |
| --- | --- | --- |
| RULE | 0.61 | $1.07 \times 10^{-40}$ |
| CGC | 0.62 | $3.34 \times 10^{-70}$ |
| HCD | 0.59 | $3.92 \times 10^{-42}$ |
| CTAT | 0.62 | $2.43 \times 10^{-52}$ |

**Table 2.**
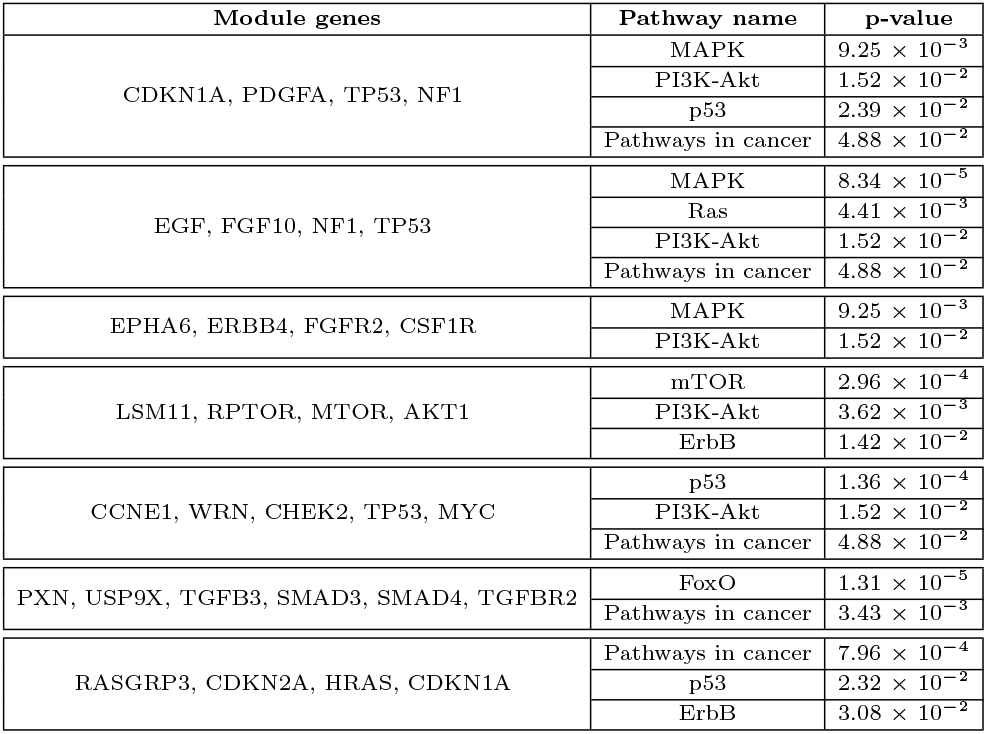
A set of obtained important modules for ovarian cancer.

In the second step, we used MG algorithm as a heuristic approach for identifying densely connected modules within a biological network (8). MG initializes by selecting a seed node based on its LS score and iteratively expands the module by adding nodes that increase the average LS score. The algorithm terminates when no further improvement is possible or when a predefined size constraint is met. The final output consists of a set of modules, each representing a potential functional unit likely to contain key driver genes and prioritized for further biological investigation. Figure 1 illustrates the workflow of the proposed two-step technique.

**Fig. 1.**
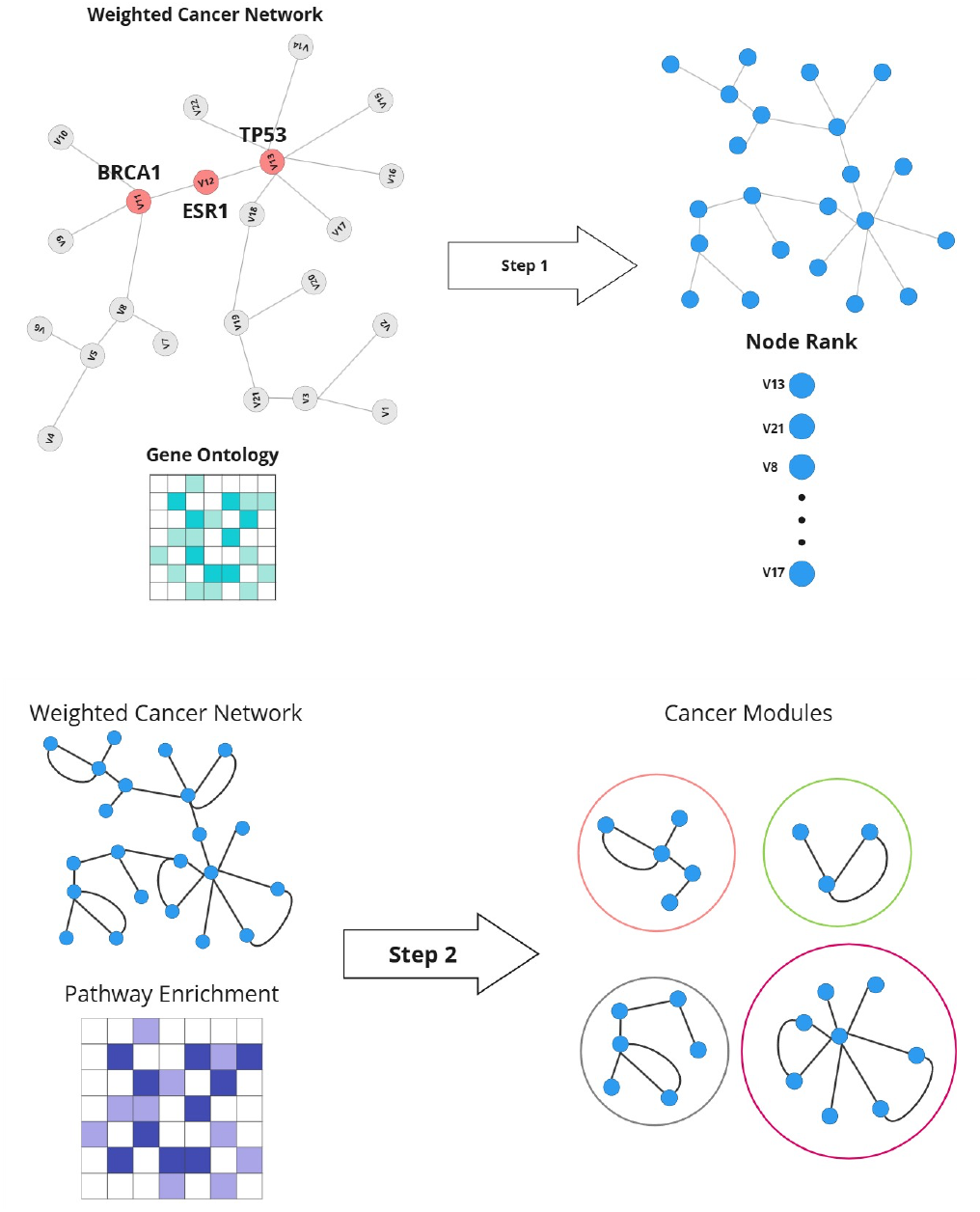
The schematic of the proposed framework: The first step involves detecting and ranking important genes, and the second step focuses on identifying informative modules.

## Results

In this study we presented two key findings. First, we evaluated the performance of our selected high-score genes by comparing them to four benchmark datasets. Second, we assessed the proposed functional modules using pathway enrichment analysis and calculated a p-value to determine their significance in cancer-related pathways.

### Evaluation of high-score selected genes

For the first evaluation part, we used AUC-ROC and the Mann-Whitney U Test (11; 12). To calculate the AUC-ROC, we identified the top *N* genes from the ranked list, where *N* equals the size of the benchmark gene set. A binary classification was assigned to each gene: 1 for genes within the top *N* (predicted as significant with high score) and 0 otherwise. The benchmark gene set served as the ground truth, with genes labeled 1 for significance and 0 for non-significance. The AUC-ROC was computed by comparing these two binary classifications, evaluating the model’s ability to distinguish between important and non-important genes. The Mann-Whitney U test compares the ranks of benchmark genes to non-benchmark genes within the high score ranked gene list. Benchmark gene ranks indicate their position in the list, while other ranks represent the positions of non-benchmark genes. The test evaluates whether benchmark genes are consistently ranked higher than non-benchmark genes and a low p-value indicates significant ranking differences, suggesting that the model effectively prioritizes benchmark genes. Table 1 shows the evaluation results for AUC-ROC and the Mann-Whitney U Test for ovarian cancer as a showcase, along with each benchmark. The reason the AUC-ROC p-value is not very low is that the benchmarks mentioned above aimed to select a large number of cancer-related genes as essential, trying to be as general as possible. As a result, comparing our rankings with theirs may not provide specific insights. However, since these benchmarks are the only relevant ones available, we decided to compare our results with them. The results for other cancer types are reported in Supplementary Table 3.

### Evaluation of modules

To ensure biological relevance and minimize redundancy, identified modules were refined through a series of steps. Modules were constrained to include 4–10 genes, striking a balance between specificity and comprehensiveness (13). Overlapping modules were compared based on their composite scores, retaining only the highest-scoring ones. Finally, quality assessments removed redundant subsets of larger modules, ensuring distinctiveness and clarity.

The evaluation process assessed each module for biological significance. Genes from each module were analyzed through pathway enrichment profiling using the KEGG database (14), identifying significant associations based on p-values *≤* 0.05. For this purpose, we used Gene Set Enrichment Analysis (GSEA) method to assess the significance of predefined gene sets based on well-known cancer-related pathways (15). We calculated the p-value as follows:

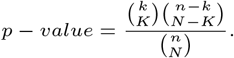

where, *N*, shows the total number of genes in the dataset for each cancer. *K* is the number of genes in a cancer-related pathway (*p*_*i*_), *n* is the number of driver genes in the detected module (*m*) and *k* is the number of genes shared between the driver module (*m*) and the pathway (*p*_*i*_) respectively.

We selected the predefined list of signaling pathways contains 11 signaling pathways that affect cell responses such as survival, proliferation, migration, differentiation, and apoptosis and promote different types of cancer. These pathways include the FoxO pathway, which regulates cell death, cell cycle, and metabolism; the Wnt pathway, crucial for cancer development and cell migration; and the MAPK pathway, involved in cell survival and proliferation. The p53 pathway controls the cell cycle and apoptosis, often altered in cancers. Estrogen pathway Regulates tissue growth and development, and is linked to some cancers. Pathways in cancer, encompasses various signaling pathways involved in cancer progression. Pathways like Ras, ErbB, and PI3K-Akt govern key cellular functions and are frequently mutated in cancers. The VEGF pathway supports tumor growth through angiogenesis, and the mTOR pathway influences cell growth and metabolism. Together, these pathways offer a comprehensive view of cancer biology. In our evaluation in this part, a module was marked as biologically significant if it showed enrichment in at least two relevant pathways, ensuring statistical and contextual relevance in cancer analysis. Table 2 displays a part of modulated genes for ovarian cancer. The first column lists the module genes, the second column indicates the contribution of each module to the predefined pathways, and the third column shows the corresponding p-value for the module in each pathway. The complete module set for ovarian cancer and the results of this part for other 12 cancer types are reported in Supplementary Tables 5-16.

## Implementation

The GenePioneer is written in Python and consists of five tools:

- `DataLoader`: Handles the extraction and preprocessing of datasets required for network construction and analysis. This tool ensures seamless loading and mapping of genes, cases, and processes, creating a foundation for downstream analyses.
- `NetworkBuilder`: Constructs a weighted, undirected gene network by leveraging shared biological processes as edge weights. It enhances the network with topological features, including node weights, centrality metrics, and entropy.
- `NetworkAnalysis`: Conducts in-depth network analysis to rank genes and detect significant modules. It utilizes LS for feature selection and applies a heuristic module-detection algorithm to identify biologically relevant clusters, ensuring high-quality module detection.
- `GeneAnalysis`: Enables the detailed exploration of individual genes within the network. With the help of this tool, users can query genes to obtain their rank, associated modules, and topological features.
- `Evaluation`: Validates the performance of the network and modules against benchmark datasets. It employs metrics such as AUC-ROC, and Mann-Whitney U Test, alongside pathway enrichment analysis, to assess the biological relevance and accuracy of the identified genes and modules.

The `GenePioneer` package is a groundbreaking resource for cancer genomics, providing tools to rank essential genes and identify cancer-related modules across various cancer types using unsupervised network-based methods. Accessible via GitHub and PyPi, the toolkit is designed with user-friendliness in mind, enabling researchers and clinicians to gain actionable insights without requiring extensive computational expertise.

## Conclusion

The `GenePioneer` toolkit was developed as a fast and straightforward way to integrate gene ranking and module detection into a practical, Python-based tool for cancer researchers. It requires minimal input, delivers clear output, and can be run within a Python environment, making it highly user-friendly and accessible to non expert programmers while supporting large-scale dataset analysis. By evaluating gene importance and identifying gene interactions within cancer networks, `GenePioneer` provides critical insights into the genetic drivers of cancer. Key features include ranking genes by their network significance and identifying the modules they belong to, which helps explore cancer-related pathways and aids in developing precise therapies. Available on GitHub and PyPi, GenePioneer’s user-centric design ensures that researchers of all skill levels can make use of its capabilities. By combining comprehensive data integration, advanced network-based analysis, and statistical rigor, `GenePioneer` stands as a versatile and impactful resource for cancer research across multiple cancer types.

## Data availability

The GenePioneer source code is freely available at https://github.com/Golnazthr/ModuleDetection

## Author contributions statement

G.T. and A.H. conceived the experiments, A.H. conducted the experiments, G.T. and A.H. analysed the results. G.T. wrote and reviewed the manuscript.

